## Supplementary Figures for "β-sheet stabilization of the island domain underlies ligand-induced LRR-RP activation of plant immune signaling"

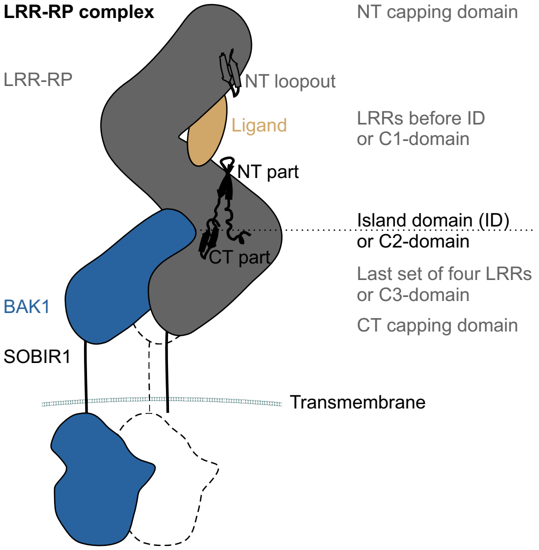


**Supplementary Fig. 1: Graphical representation of the LRR-RP – ligand – BAK1 – SOBIR1 receptor complex.** Overview of the LRR-RP nomenclature.

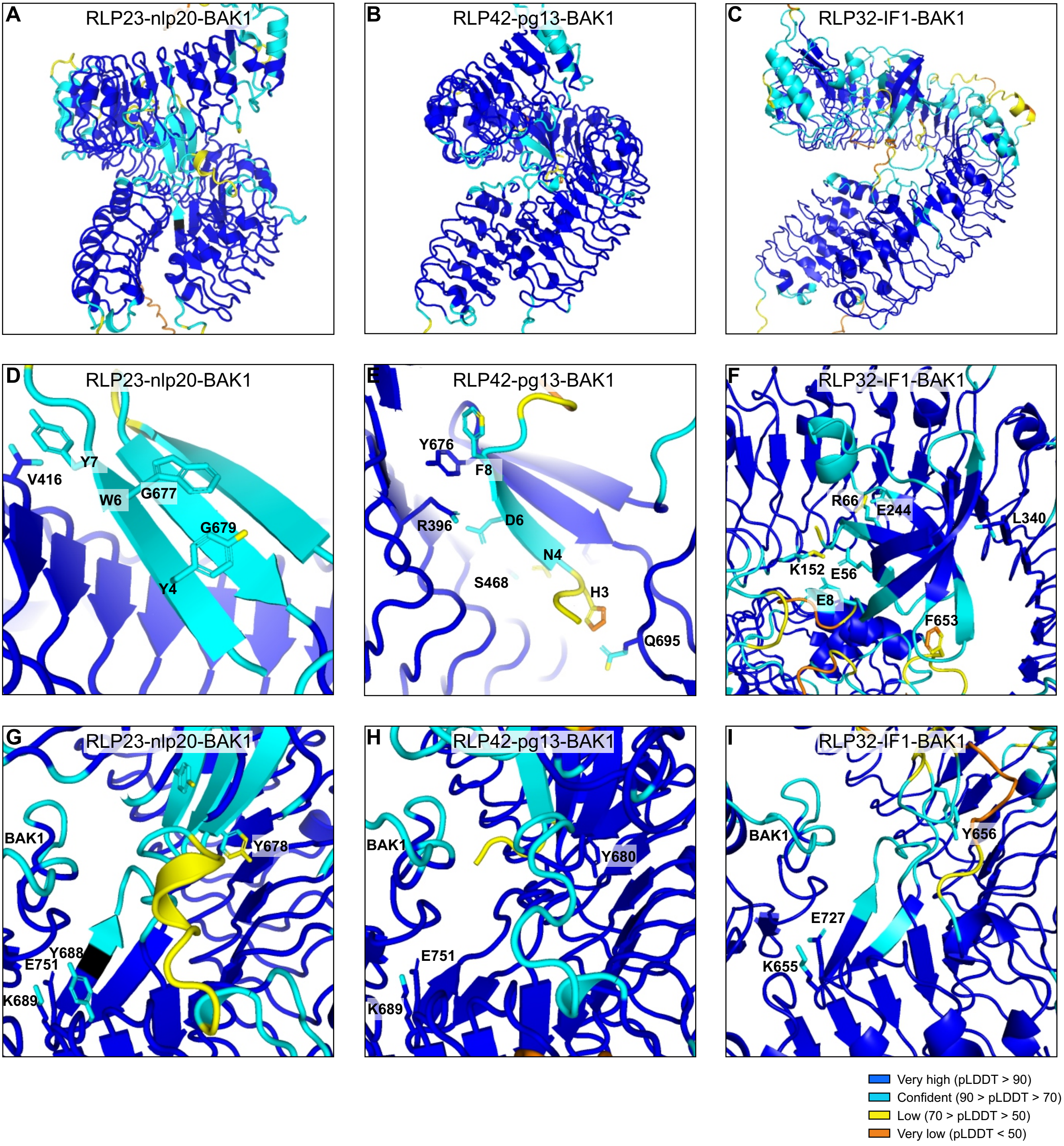


**Supplementary Fig. 2: Visualization of the predicted local distance difference test (pLDDT) score for the AF3 predictions of RLP23 (left), RLP42 (middle) and RLP32 (right).**
pLDDT is a per-atom confidence estimate on a 0-100 scale where a higher value indicates higher confidence and usually a more accurate prediction, here depicted for the LRR-RP complex predictions using the pymol extension pymol-color-alphafold, coloring as per legend. **A-C)** Overview of the complete receptor complex prediction. **D-F)** Zoom in on the ligand-binding interface. **G-I)** Zoom in on the interaction interface between the NT part of the ID and BAK1. Pdb files can be found in Supplementary Dataset 1.

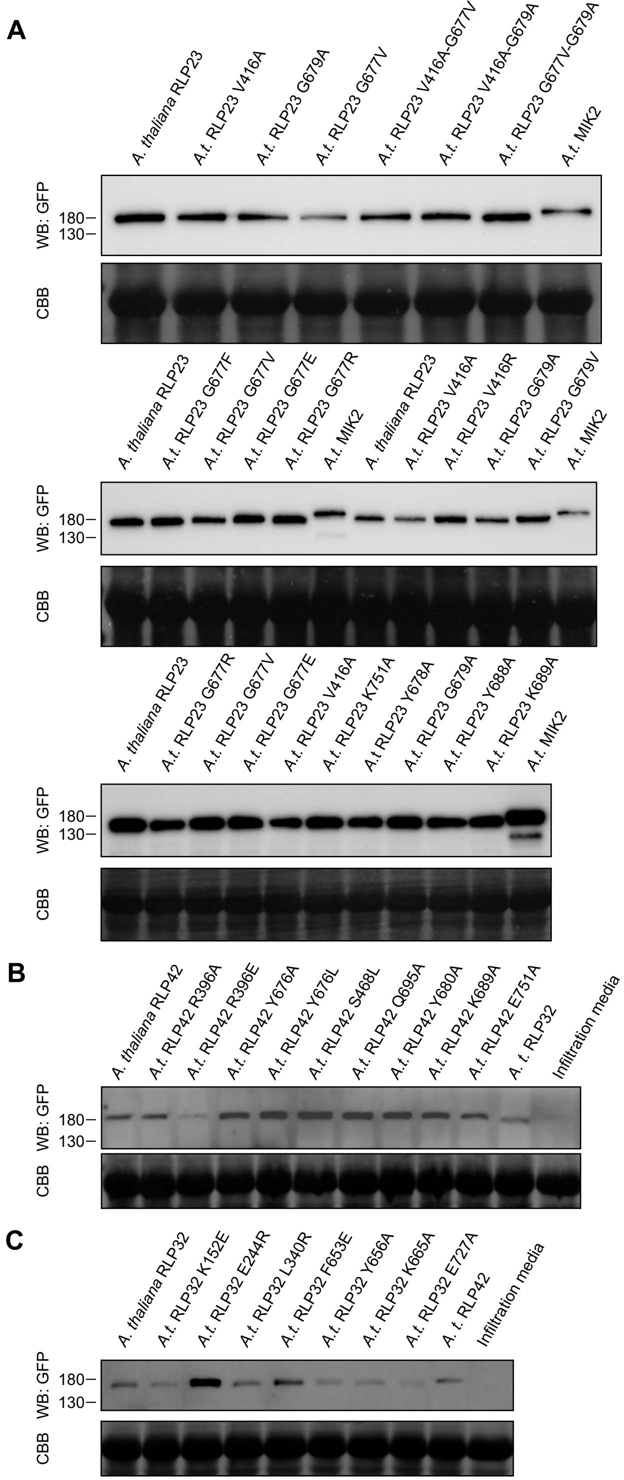


**Supplementary Fig. 3: Western blotting following heterologous expression of the LRR-RPs and their respective variants in *N. benthamiana*.**

**A-C)** Western blot 72 h post-Agrobacterium infiltration of respectively RLP23, RLP42 and RLP32. The western blots were probed with α-GFP (B-2) HRP as the receptor had a C-terminal GFP tag (top) and subsequently stained with CBB as a loading control (bottom). MIK2, RLP32 and RLP42 were used as positive control for heterologous expression respectively in A, B and C, as they share the same expression vector as the receptor (variants) of interest.

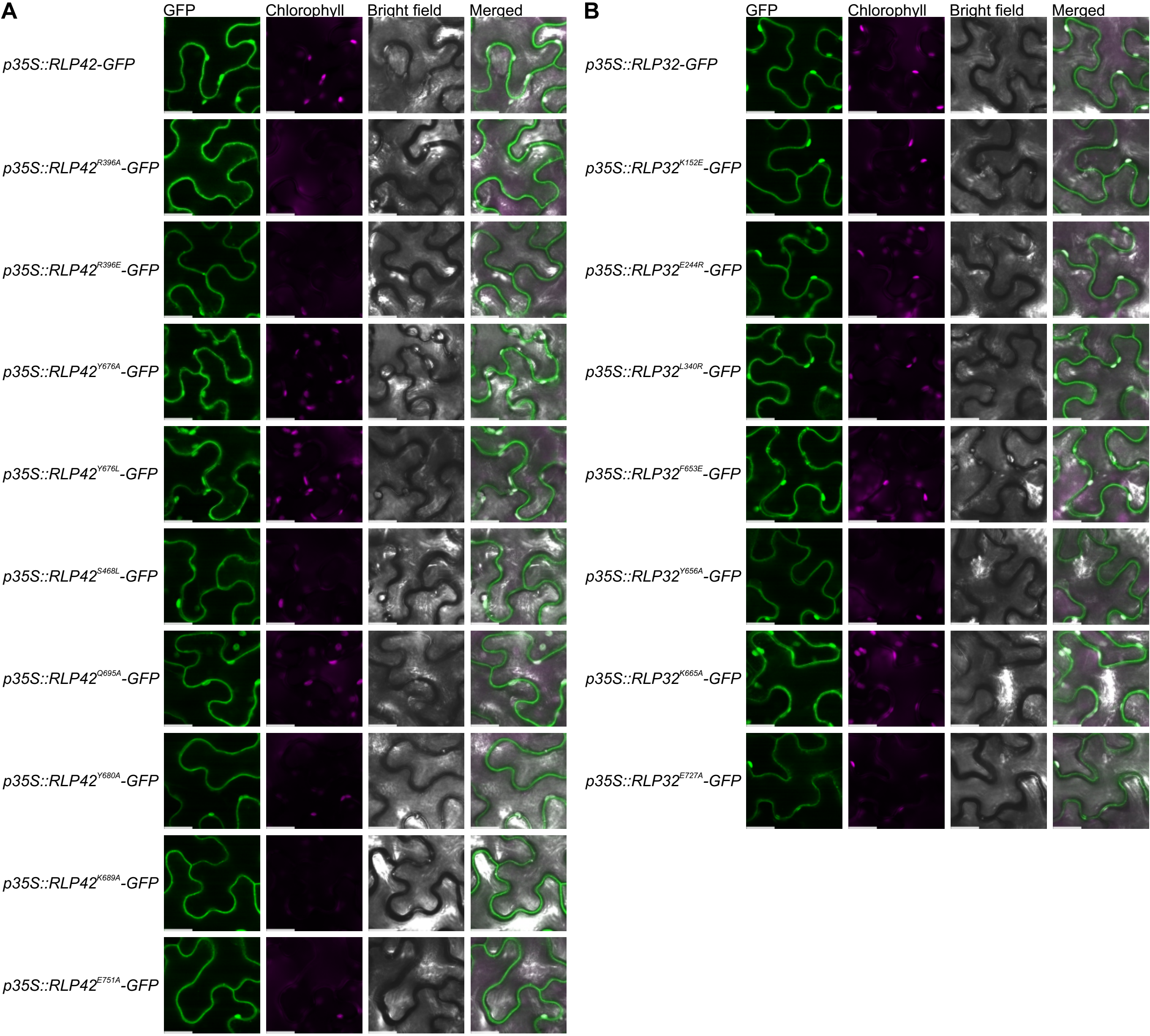


**Supplementary Fig. 4: Confocal microscopy following heterologous expression of the LRR-RPs and their respective variants in *N. benthamiana*.**
**A-B)** Confocal microscopy (GFP, Chlorophyll B and Bright Field) following Agrobacterium infiltration of respectively RLP42 (A) and RLP32 (B) and their respective variants (72 h). All confocal microscopy images were taken with the same image settings and identically modified. White scale bar represents 20 µm.

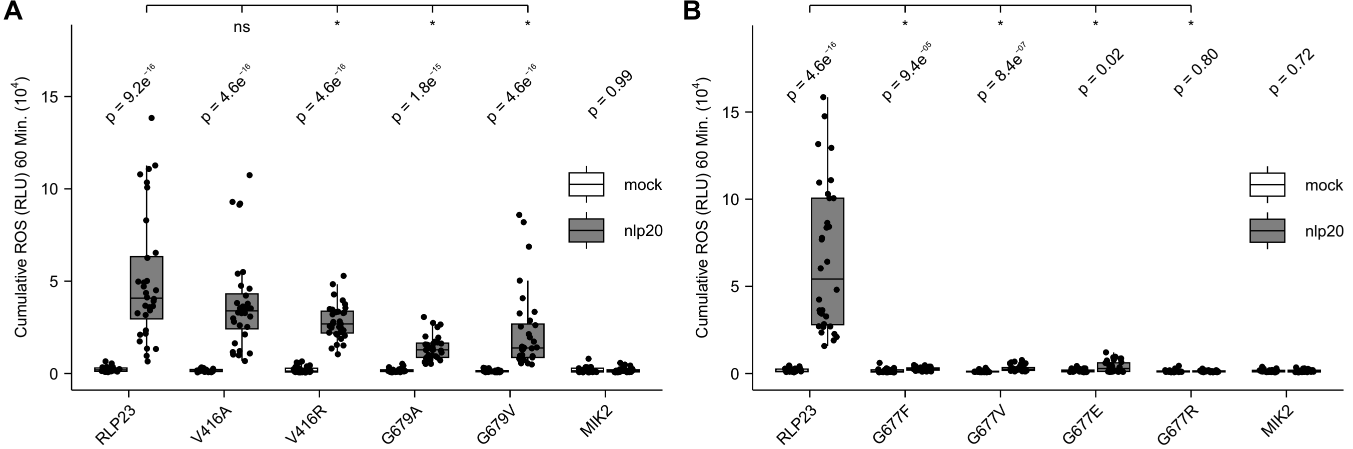


**Supplementary Fig. 5: Single AA changes to diverse residues within the predicted ligand-binding interface affect the functionality of the RLP23. A-B)** ROS production (4 to 60 min) in cumulative RLUs post treatment with $H_2O$ (white) or 1 µM nlp20 (gray). Eight independent biological replicates (n = 8 plants) were performed, with each biological replicate represented by at least three technical replicates. Significance was tested by performing non-parametric Wilcoxon-Mann-Whitney tests between both mock and ligand RLP23 (variants) as well as ligand-treated RLP23 vs specific variants. The asterisks indicate a significant difference of $p < 0.05$.

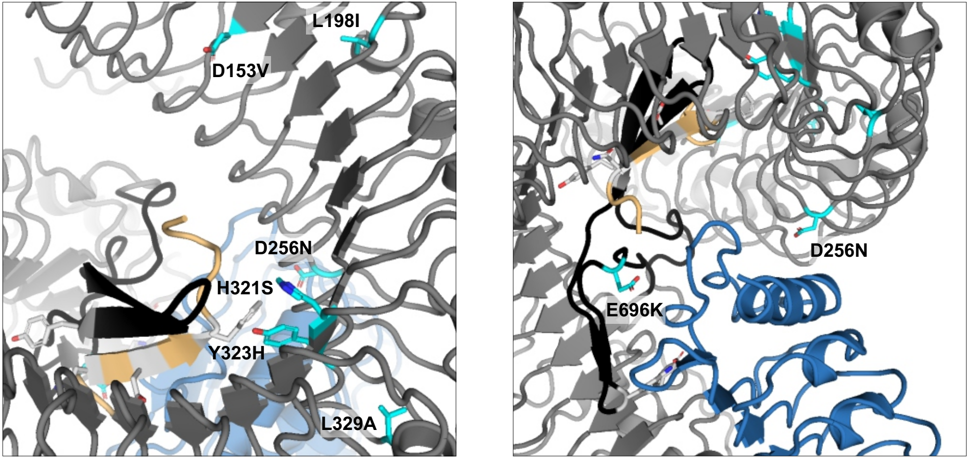


**Supplementary Fig. 6: Earlier characterized single AA changes affect ligand binding and BAK1 recruitment by RLP42.** Structural representations of the tripartite complexes of RLP42-pg13-BAK1. The ID is highlighted in black, other LRR-RP domains in dark grey. The ligand is depicted in yellow, and BAK1 in blue. Residues highlighted in light blue were earlier shown to affect RLP42 functionality [9]. Pdb files can be found in Supplementary Dataset 1.
